# Integrated computational and *in vivo* models reveal Key Insights into Macrophage Behavior during bone healing

**DOI:** 10.1101/2021.12.09.471980

**Authors:** Etienne Baratchart, Chen Hao Lo, Conor C. Lynch, David Basanta

## Abstract

Myeloid-derived monocyte and macrophages are key cells in the bone that contribute to remodeling and injury repair. However, their temporal polarization status and control of bone-resorbing osteoclasts and bone-forming osteoblasts responses is largely unknown. In this study, we focused on two aspects of monocyte/macrophage dynamics and polarization states over time: 1) the injury-triggered pro- and anti-inflammatory monocytes/macrophages temporal profiles, 2) the contributions of pro-versus anti-inflammatory monocytes/macrophages in coordinating healing response. Bone healing is a complex multicellular dynamic process. While traditional *in vitro* and *in vivo* experimentation may capture the behavior of select populations with high resolution, they cannot simultaneously track the behavior of multiple populations. To address this, we have used an integrated a coupled ordinary differential equations (ODEs)-based framework describing multiple cellular species to *in vivo* bone injury data in order to identify and test various hypotheses regarding bone cell populations dynamics. Our approach allowed us to infer several biological insights including, but not limited to,: 1) anti-inflammatory macrophages are key for early osteoclast inhibition and pro-inflammatory macrophage suppression, 2) pro-inflammatory macrophages are involved in osteoclast bone resorptive activity, whereas osteoblasts promote osteoclast differentiation, 3) Pro-inflammatory monocytes/macrophages rise during two expansion waves, which can be explained by the anti-inflammatory macrophages-mediated inhibition phase between the two waves. In addition, we further tested the robustness of the mathematical model by comparing simulation results to an independent experimental dataset. Taken together, this novel comprehensive mathematical framework allowed us to identify biological mechanisms that best recapitulate bone injury data and that explain the coupled cellular population dynamics involved in the process. Furthermore, our hypothesis testing methodology could be used in other contexts to decipher mechanisms in complex multicellular processes.

**Author Summary:** Myeloid-derived monocytes/macrophages are key cells for bone remodeling and injury repair. However, their temporal polarization status and control of bone-resorbing osteoclasts and bone-forming osteoblasts responses is largely unknown. In this study, we focused on two aspects of monocyte/macrophage population dynamics: 1) the injury-triggered pro- and anti-inflammatory monocytes/macrophages temporal profiles, 2) the contributions of pro-versus anti-inflammatory monocytes/macrophages in coordinating healing response. In order to test various hypotheses regarding bone cell populations dynamics, we have integrated a coupled ordinary differential equations-based framework describing multiple cellular species to *in vivo* bone injury data. Our approach allowed us to infer several biological insights including: 1) anti-inflammatory macrophages are key for early osteoclast inhibition and pro-inflammatory macrophage suppression, 2) pro-inflammatory macrophages are involved in osteoclast bone resorptive activity, whereas osteoblasts promote osteoclast differentiation, 3) Pro-inflammatory monocytes/macrophages rise during two expansion waves, which can be explained by the anti-inflammatory macrophages-mediated inhibition phase between the two waves. Taken together, this mathematical framework allowed us to identify biological mechanisms that recapitulate bone injury data and that explain the coupled cellular population dynamics involved in the process.

## INTRODUCTION

The tightly-coupled relationship between bone-forming osteoblasts and bone-resorbing osteoclasts in bone remodeling and healing is well established^1^. Bone remodeling is initiated by osteoclastic turnover of aged and compromised bone tissue. Molecular cues derived from bone resorption subsequently drive mesenchymal precursor expansion and differentiation into osteoblasts for formation of new bone^1^. Bone healing on the other hand begins with osteoblastic bone callus deposition that is subsequently remodeled by osteoclasts^1^. Beyond this classic paradigm of the bone modeling unit (BMU), studies are increasingly identifying other cellular populations and factors that also contribute to the maintenance of bone. Macrophages of the myeloid lineage play critical roles in inflammation, wound healing and cancer progression^2^. Recent studies have also shed light on their contribution to bone biology. While osteoclasts have traditionally been known as the tissue resident macrophage of the bone, more recent studies identified a novel population of bone-resident macrophages, osteomacs, which facilitate osteoblast bone formation^3, 4^. Additionally, in the context of bone healing, macrophages have been documented to rapidly infiltrate sites of bone injury to clear cellular debris in a process called efferocytosis and elicit subsequent inflammatory response and mineralized callus formation^1^. Monocytes and macrophages are major components of the bone immune infiltrate following injury^1, 5–7^. Recent studies using genetic or pharmacological depletion of macrophages demonstrated significantly delayed time to bone repair ^4, 5, 8–10^. The diversity of macrophage function owes to its versatility in polarizing and responding to environmental cues^1, 6–8^. These critical functions ensure the right temporal sequence of events necessary for healthy and timely bone repair after injury. For instance, IL-4 and TNFα have been shown to promote different macrophage polarization states and impact bone healing ^11–13^. As an example, acute pro-inflammatory factors such as TNFα can improve bone repair while prolonged administration has the opposite effect^11, 12^. There are, however, a number of gaps in our understanding of monocyte and macrophage population and polarization behavior, including but not limited to: 1) the contributions of pro-versus anti-inflammatory macrophages in coordinating bone injury response, 2) whether macrophages are directly involved in control of osteoclasts and osteoblasts populations and activities during bone injury, and 3) the main mechanisms that govern pro- and anti-inflammatory macrophages population dynamics.

While *in vitro* and *in vivo* experimentation techniques can capture the behavior of individual populations with high resolution, they do not allow for understanding the simultaneous interplay between multiple cell types whose numbers change over time. This obstacle can be overcome with the integration of experimental data to computational approaches in order to model the interactions occurring during bone injury repair. This type of approach has already been applied to other disease contexts like cancer^14–23^. Amongst the possible types of modeling approaches, agent-based models, such as discrete-continuum Hybrid Cellular Automata can examine mechanisms at the cellular scale leading to emergence of non-trivial macroscopic patterns^24^. One advantage of such an approach, is the possibility to inform the model with experimentally measured parameters. However, these parameters, such as macrophage polarization rate for example, can sometimes be exceptionally difficult to measure *in vivo* or *in vitro*. On the other hand, systems of Ordinary Differential Equations (ODEs) model individual populations over time under a well-mixed assumption and are often used to estimate *in vivo* parameters. While they do not describe cellular mechanisms as finely as agent-based models, their relative computational simplicity make them a convenient tool to identify key parameters through data fitting^25–27^. Multiple mathematical approaches have been used to study bone remodeling and repair ^28–32^. The vast majority use systems of ODEs to model bone cell populations in homeostatic bone remodeling and bone disease such as osteoporosis and multiple myeloma^33–38^. Bone remodeling is a physiological program that is tightly regulated spatially and temporally. Other groups have considered the role of space in the process and represented cell population either as continuous spatial field, describing the dynamics by a set of partial differential equations (PDE) ^39, 40^, or as individual agents by an agent-based model approach ^23^. These models have largely focused on the interaction between bone-building osteoblasts and bone-resorbing osteoclasts, mostly ignoring the role of immune and inflammatory cells. Although these models have addressed biologically and clinically relevant questions, very few studies, one of which is from our group^41^ have quantitatively compared predictions of bone injury dynamics to longitudinal biological data. Some studies have included the role of inflammatory cells like macrophages, but they remain theoretical and have not been experimentally validated^42, 43^. The role of inflammation, and that of macrophages in particular, is recognized as being key for coordinating the bone injury response *in vivo* but, to date, how monocyte/macrophage populations coordinate it and interact directly with osteoblasts and osteoclasts (and vice versa) during bone remodeling has not been thoroughly examined^1, 6^. Here we use experimental, in combination with published data, to integrate osteoblasts, osteoclasts, bone, naïve, pro- and anti-inflammatory monocytes and macrophages into a coupled ODE model of the bone ecosystem. This approach allowed for the interrogation of key hypotheses that explain the bone healing program, such as the polarization and clearance dynamics of monocytes/macrophages, interactions between anti-inflammatory macrophages and pro-inflammatory monocytes/macrophages, and how pro- and anti-inflammatory monocytes/macrophages modulate osteoclast and osteoblast behaviors. We posit this integrated approach can be used to uncover mechanisms driving bone injury repair dynamics and to identify key strategies aimed at shortening bone healing times.

## RESULTS

### Quantitative data of cell populations during bone injury repair dynamics

During non-critical bone injury healing, the following sequence of steps occur: early inflammation and hematoma formation, direct intramembranous bone deposition into a mineralized callus by osteoblasts^44–46^, and callus remodeling by osteoclasts^1, 3, 9, 47^. Bone-forming osteoblasts and bone-resorbing osteoclasts are critical mediators of these steps, and their numbers shift accordingly during each phase of repair. In order to temporally quantitate bone cells during injury, we extracted multi-cellular longitudinal data from an experimental model of bone injury repair whereby non-critical epiphyseal fracture was generated in mice by direct intratibial injection^41, 47–50^ (Fig. 1a). In the injured tibias of the mice, bone volume and cell populations numbers were quantified at baseline (day 0) and at day 1, 2, 3, 7 and 14 (n = 5/time point) following injury. High-resolution μCT analysis of the site of bone injury demonstrated changes in bone volume (BV/TV) subsequent to bone injury (Fig. 1b). Bone volume remained diminished over a 48-hour period prior to a five-day long expansion, beyond baseline levels. By day 14, the bone volume returned toward homeostasis. Consistent with other published observations, the overall bone volume dynamics were accompanied by corresponding sequential waves of osteoblast and osteoclast numbers^51 50 52^ (Fig. 1b). Interestingly, the overlaid data reveal alternating waves of osteoblasts and osteoclasts. In the same longitudinal study, the contralateral tibia from each mouse was additionally subject to flow cytometry to derive dynamics of total and polarized myeloid populations ^2, 4, 5, 7, 53–107^ (Fig. 1b). The myeloid dataset shows that pro-inflammatory monocytes and macrophages spiked within the first 48 hours while anti-inflammatory macrophages were observed between 24 and 72 hours (Fig. 1b). Importantly, a fainter but prolonged secondary wave of pro-inflammatory monocytes-macrophages was noted, an observation which is in line with past studies in other inflammatory contexts^52, 108–110^.

**Fig 1.**
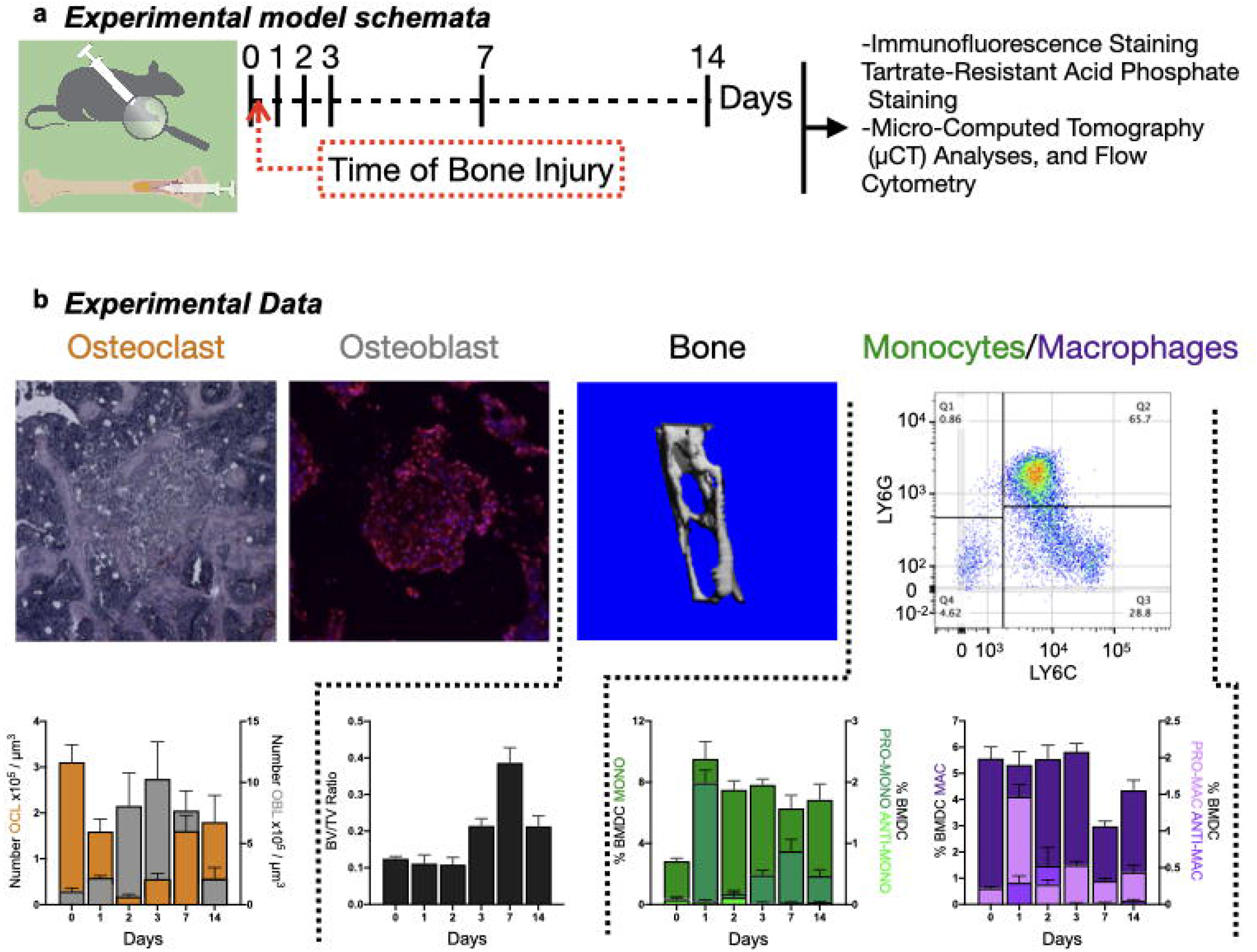
Experimental quantification of osteoblast, osteoclast, bone volume, and monocyte-macrophage over time during bone injury. **a** schematic summarizing the experimental system and the time course and different measurements performed. **b** from left to right: Decalcified bones were stained and quantified for OCL by tartrate-resistant acid phosphatase (TRAcP) staining (top left panel; red). Temporal quantification of OCL population was then assessed (bottom left panel); Decalcified bones were stained and quantified for OBL by RUNX2 immunofluorescence staining (second top panel; red). Temporal quantification of OBL population was then assessed (bottom left panel); micro-computed tomography revealed trabecular bone status. Representative images (third top panel) and corresponding quantitative analyses of bone volume on the top panel (BONE; BV/TV, second bottom panel). Flow cytometry was to gate and quantify monocytes-macrophages by the use of CD11b, Ly6C, Ly6G markers (Top right panel). Temporal quantification of naïve, pro-inflammatory and anti-inflammatory monocytes-macrophages populations was then assessed (bottom right panel).

### Mathematical modeling reveals key insights into myeloid behaviors during bone injury repair

In order to shed light on monocyte-macrophage dynamics, we interrogated hypotheses regarding population dynamics, differentiation, lifespan and plasticity. To this end we built a coupled ordinary differential equations-based framework describing seven cell populations as well as the bone volume temporal dynamics. The cell populations we considered in the model were bone-building osteoblasts, bone-resorbing osteoclasts, naïve monocytes, pro-inflammatory monocytes, naïve macrophages, pro and anti-inflammatory macrophages. To properly integrate cell population temporal data into this framework, we first curated common literature observations and hypotheses regarding osteoblasts, osteoclasts, monocyte and macrophage behavior during tissue injury healing (Table 1). In the model, the initial dynamics (osteoblast expansion, osteoclast decrease, macrophage polarization, monocyte infiltration) are triggered by injury factors^108^. We assume that the amount of factors released from an injury are proportional to the bone damage induced and are the primary driver of myeloid response. Myeloid cells are known to infiltrate the bone and polarize into pro-inflammatory status to clear cellular debris when exposed to injury-associated factors (Table 1)^1^. Therefore, an injury variable was included in the model that drives the initial pro-inflammatory response by monocytes/macrophages. In the model, this injury variable is being depleted by a decay rate term that is proportional to the number of pro-inflammatory cells (Equations on Supplemental Fig. 1 and Supplemental Fig. 3)

Browsing existing literature (Table 1), we identified various hypotheses regarding monocyte/macrophage control of osteoclast and osteoblast numbers, and the mechanistic relationship between pro- and anti-inflammatory myeloid cells (Fig. 2 and Table 1). These hypotheses pertain to these three aspects of cell population dynamics:

a. Osteoclast dynamics

1. Pro-inflammatory monocytes/macrophages stimulate osteoclast expansion; Anti-inflammatory monocytes/macrophages inhibit osteoclast formation and life span
2. Pro-inflammatory monocytes/macrophages stimulate osteoclast expansion; Osteoblasts inhibit osteoclast formation and life span
3. Osteoblasts stimulate osteoclast expansion; Anti-inflammatory macrophages inhibit osteoclast formation and life span
b. Osteoblast dynamics

1. Anti-inflammatory factors stimulate osteoblast expansion
2. Injury factors stimulate osteoblast expansion
c. Monocyte-Macrophage dynamics

1. Injury factors drive both pro-inflammatory monocytes/macrophages and anti-inflammatory macrophages polarization; Anti-inflammatory macrophages suppress pro-inflammatory macrophages
2. Injury factors drive pro-inflammatory monocytes/macrophages polarization; Pro-inflammatory monocytes/macrophages drive anti-inflammatory macrophages polarization; Anti-inflammatory macrophages suppress pro-inflammatory macrophages
3. Injury factors drive resident pro-inflammatory macrophages polarization; pro-inflammatory macrophages repolarize into anti-inflammatory macrophages when phagocytosing cellular debris; Anti-inflammatory macrophages naturally repolarize into pro-inflammatory macrophages in absence of stimulus (plasticity); Pro-inflammatory macrophages drive pro-inflammatory monocyte polarization.

**Fig 2.**
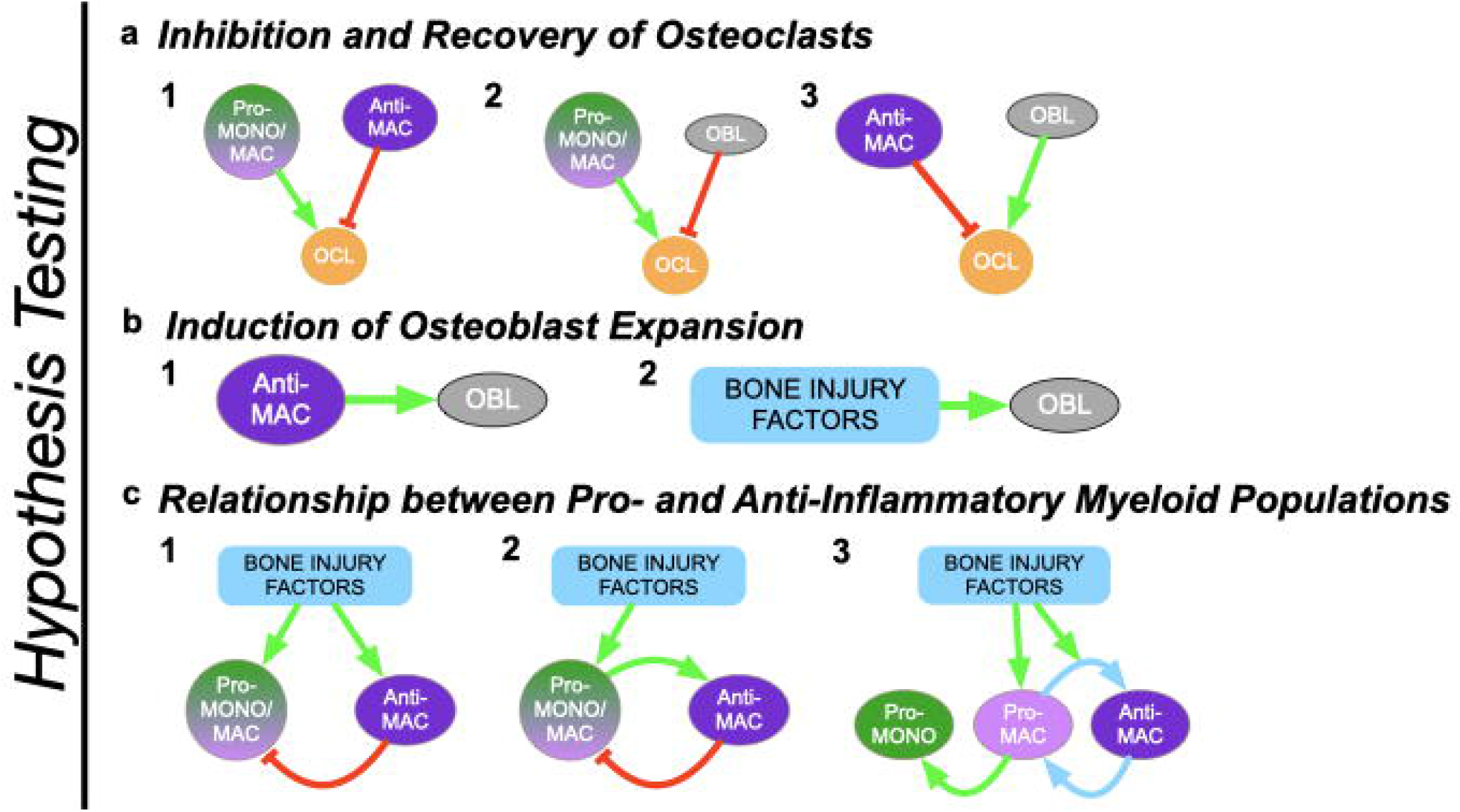
Comprehensive combinatorial modeling pipeline is built to identify relevant myeloid behaviors necessary to recapitulate *in vivo* bone injury repair data. Literature curation reveals sets of well-established competing biological mechanisms potentially governing modulation of osteoclast formation (**a**), regulation of osteoblast formation (**b**), and relationships between pro- and anti-inflammatory myeloid cells (**c**). Model adopts all combinations of hypotheses regarding these mechanisms to recapitulate *in vivo* data. Comparing model fits resulting from each hypothesis combination reveals best-fitting models.

Whereas evidence for all these mechanisms have been shown *in vitro* or *in vivo*, our goal here is to identify the ones that can recapitulate *in vivo* dynamics, in order to define main mechanisms that drive bone injury dynamics. We focus here on parsimonious hypothesis combinations, where for a given cell dynamics aspect (a, b or c), one mechanism only is considered, e.g c1 as opposed to c1&2.

We described the hypotheses using ordinary differential equations and integrated them into a mathematical model framework to assess the ability of each hypothesis combination to recapitulate the experimental data. This resulted in 18 ODE models, each describing a unique combination of hypotheses. In each permutation, we assessed how well the model fitted to the experimental data (Table 2). The models were ranked based on their goodness of fit, which was measured by the Akaike Information Criterion (AIC, Table 2), and number of residuals lower than 1 (Table 5). The fits to experimental data were obtained in two different ways, with the choice of two different functionals to minimize, *J*_2_ and *J*_∞_. Here we present the results obtained with *J*_∞_ but the conclusions remained the same with *J_2_*. Results indicate hypotheses combination *a3-b2-c2* (AIC of 39, Fig. 3, equations presented in Supplementary Fig.1) shows the best fit to experimental data. The second, third and fourth best fits were obtained by *a3-b1-c2* (AIC of 42, Supplementary Fig. 2), then a3-b1-c1 (AIC of 45, Supplementary Fig. 4) and a3-b2-c1 (AIC of 46, Supplementary Fig. 5). The AICs of the remaining combinations were substantially lower (Table 2). Looking to another metric of goodness of fit, the number of residuals lower than 1, *a3-b2-c2* is clearly the best combination, with 25 residuals lower than one, and the rest of the models is far apart. With 15, 13 and 14 residuals lower than one, respectively, *a3-b1-c2*, a3-b1-c1 and a3-b2-c1 rank pretty low regarding to the residuals metric (Table 5). Interestingly, some hypothesis combinations do better than *a3-b1-c2*, a3-b2-c1 and a3-b1-c1 in term of number of residuals lower than 1, but worst in term of AIC. In conclusion, combination *a3-b2-c2* does substantially better than all other combinations for both goodness of fit criteria (and for both *J*_2_ and *J*_∞_ optimizations). Of note, the best-fitting hypothesis *a3-b2-c2* (Fig. 3) assumes osteoblasts are the main osteoclastogenesis driver (Fig. 2*a3*), and that anti-inflammatory macrophages play an important role in suppressing osteoclasts and pro-inflammatory macrophages^93^ (Fig. 2*c2*). It also suggests that initial osteoblast expansion is driven by factors associated with the onset of bone injury^108^. The coupled mathematical model also allows for the estimation of polarization rates over time for pro-and anti-inflammatory macrophages in these two scenarios (Table 3). By comparison, some hypotheses combinations such as *a2-b2-c3* yielded significantly poorer fits (AIC of 113 for *a2-b2-c3*, the worst fitting one, Table 2 and Fig. 4, AIC of 79 for *a2-b1-c1*, the second worst fitting one, Table 2 and Supplementary Fig. 6). This last result demonstrates that some cellular mechanisms and behaviors, though well-established in orthopedics (e.g. osteoclast stimulation by pro-inflammatory monocytes/macrophages; osteoclasts inhibition by osteoblasts through signals like OPG) are not able to recapitulate experimental observations in specific physiological contexts. Taken together, these results show, through our integrative hypothesis combination testing framework, the minimal set of cell-cell behaviors necessary to recapitulate bone injury temporal dynamics.

**Fig 3.**
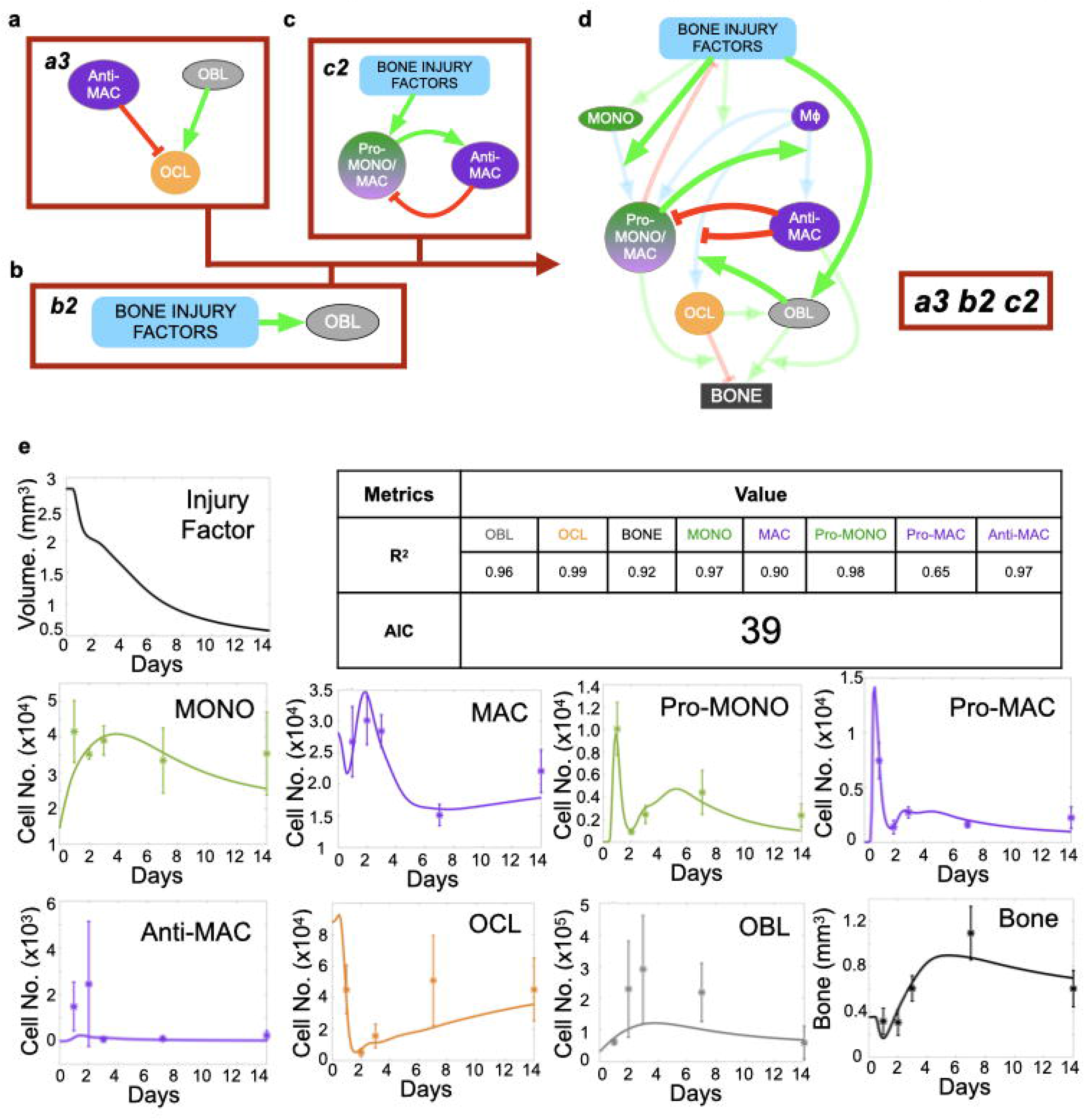
The best fitting hypothesis combination model integrates hypotheses *a3, b2* and *c2* (red boxes in **a-c**). **a** mechanism *a3* assumes that osteoblasts and anti-inflammatory macrophages promote and inhibit osteoclast formation, respectively. **b** mechanism *b2* assumes that injury factors promote osteoblast expansion. **c** mechanism *c2* assumes that injury factors promote pro-inflammatory monocytes/macrophages polarization. Pro-inflammatory monocytes/macrophages promote anti-inflammatory macrophages polarization, which in return drive depolarization of monocytes/macrophages back to the naive state. **d** schematic representation of the model using *a3-b2-c2* hypothesis combination. Arrows represent positive (green) or negative (red) types of cellular interactions. **e** Temporal plots and corresponding goodness of fit metrics (AIC and R2s) across all populations, obtained through *J*_∞_ minimization.

**Fig 4.**
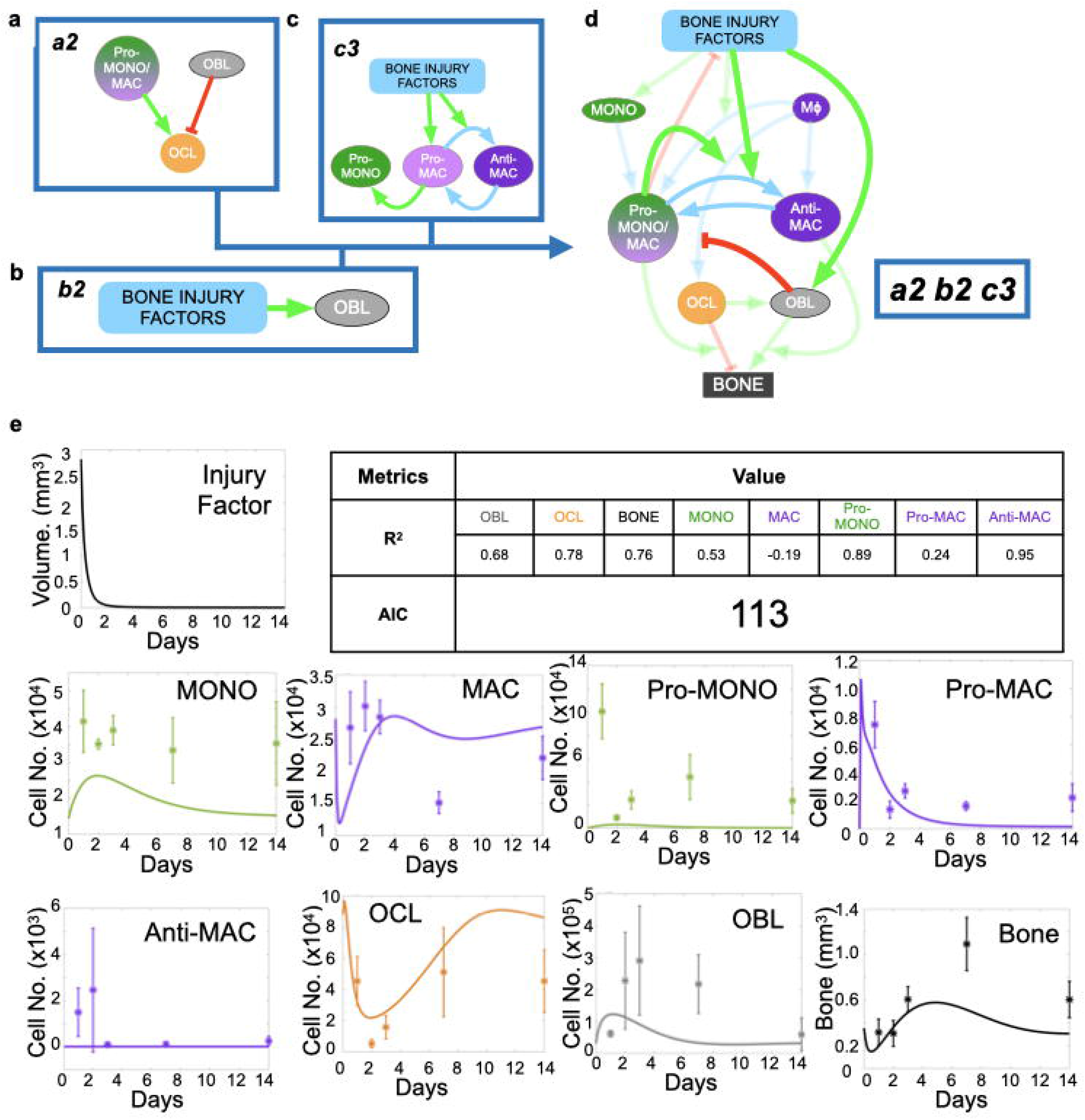
Other hypotheses combinations fail to recapitulate in vivo data. The example of the worst fitting hypothesis *a2-b2-c1* fails to recapitulate *in vivo* data. **a** mechanism *a2* assumes that pro-inflammatory monocytes/macrophages and osteoblasts promote and inhibit osteoclast formation, respectively. **b** mechanism *b2* assumes that injury factors promote osteoblast expansion. **c** mechanism *c1* assumes that injury factors promote pro-inflammatory monocytes/macrophages and anti-inflammatory macrophages polarization. Anti-inflammatory macrophages drive depolarization of monocytes/macrophages back to the naive state. **d** schematic representation of the model using *a2-b2-c1* hypothesis combination. Arrows represent positive (green) or negative (red) types of cellular interactions. **e** temporal plots and corresponding goodness of fit metrics (AIC and R2s) across all populations, obtained through *J*_∞_ minimization.

### Model simulations are consistent with independent published experimental data

Analysis of the literature reveals several factors that are important regulators of bone injury repair, such as tumor necrosis factor alpha (TNFα), interleukin-4 (IL-4), interferonγ (IFNγ) and oncostatin M (OSM) ^70, 100, 111–118^. For example, studies in mice genetically deficient for OSM exhibited reduced bone formation and osteoblasts numbers at the non-critical bone injury site^50, 119^. OSM is produced by anti-inflammatory macrophages and promotes osteoblast expansion and activity ^50, 119–121^. To assess the robustness of our bone injury repair mathematical model we simulated the effect of OSM depletion on osteoblast number and determined if the model would recapitulate the qualitative temporal dynamics of osteoblast, osteoclast population and bone volume as shown in an independent experimental dataset^50^. We found that reducing the effect of anti-inflammatory macrophages on osteoblast expansion by 50%, mineralization activity by 50% and osteoclast and osteoclast inhibition by 80% yielded similar osteoblast, osteoclast and bone dynamics to those obtained from the OSM-deficient mice (Fig. 5). Of note, osteoblast and bone levels are below baseline osteoclast number and remain largely unchanged between treatment and control in both the experimental data and model predictions. While not examined *in vivo* by the independent study, our mathematical model additionally generated corresponding predictions of the effect of OSM depletion on monocyte/macrophage dynamics (Fig. 5). Interestingly, OSM depletion increased anti-inflammatory macrophage population and transiently decreased pro-inflammatory populations. Collectively, our model predictions are in qualitative accordance with this independent experimental dataset. This suggests that the model can be used for understanding the roles of myeloid cells in the bone ecosystem during bone injury healing and for developing therapies to accelerate and improve the process.

**Fig 5.**
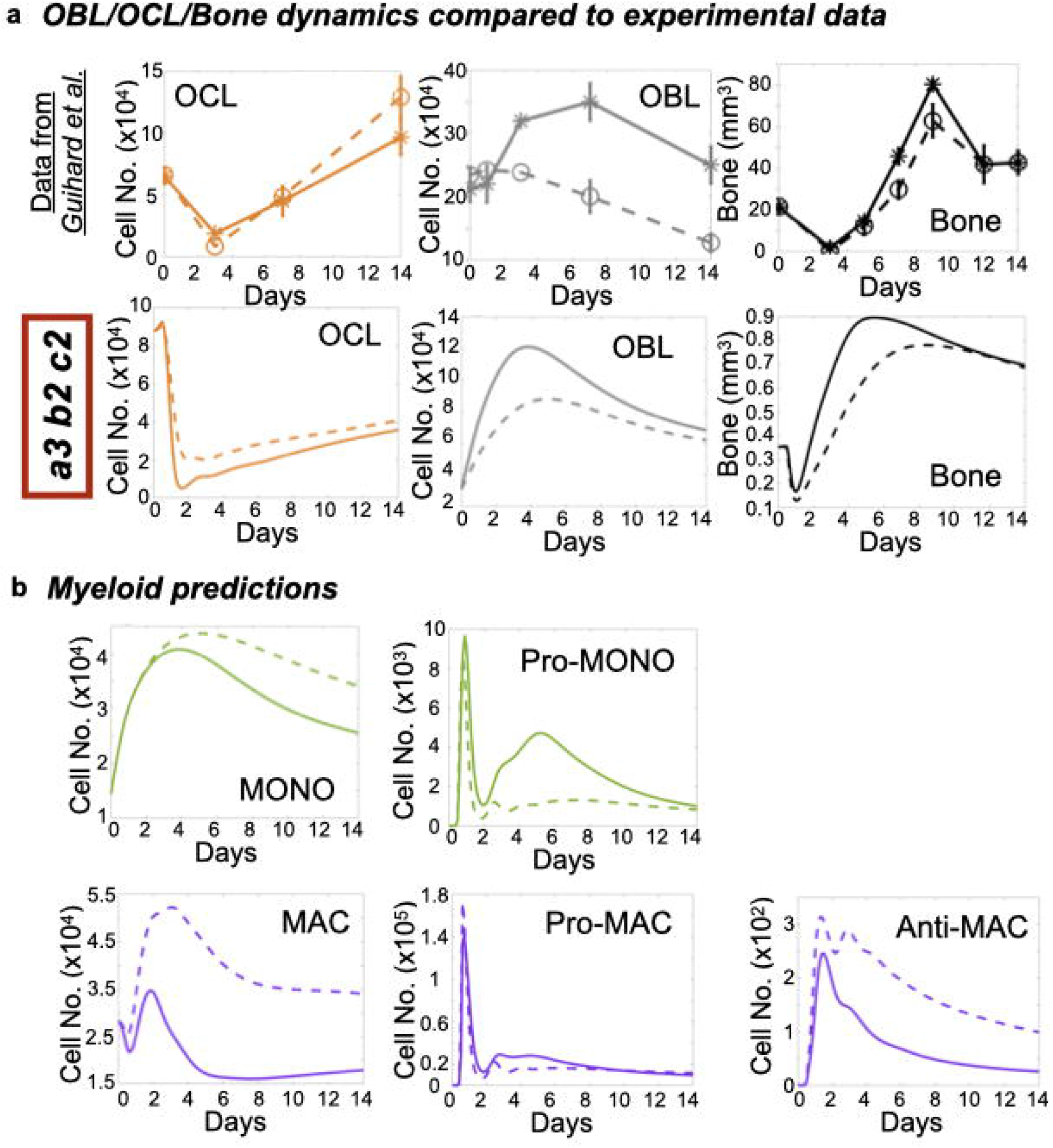
Bone repair dynamics in oncostatin M (OSM)-depleted bone predictions. **a** bone repair temporal data for OCL, OBL and bone, in presence or absence of OSM, is retrieved and plotted from a murine *in vivo* bone fracture healing study performed by *Guihard P, et al*. on the top panel (solid line = WT, dashed line = OSM-null; http://doi.org/10.1016/j.ajpath.2014.11.008). Reduction in OBL bone formation rate and mineralization activity, allow model to qualitatively reproduce OBL, OCL and bone dynamics in OSM-null dataset (lower panel; solid line = unmodulated, dashed line = OSM^-/-^). **b.** corresponding Myeloid populations predictions with reduced OBL formation rate and mineralization activity (lower panel; solid line = unmodulated, dashed line = OSM^-/-^), for which no data was available in *Guihard P, et al*. Simulations were obtained with model *a2-b2-c1*, calibrated on the injury data (Fig. 4) through *J*_∞_ minimization.

## DISCUSSION

Much remains to be discovered about how cells in the bone ecosystem collectively orchestrate the bone injury repair. Pharmacological and genetic experimental approaches can provide information as to the importance of key populations of cells, such as macrophages, but these approaches seldomly address the direct and indirect effects of other cell types involved in bone injury repair. Mathematical modeling has the advantage of being able to consider complex biological processes resulting from the interactions between several cellular populations, but their relevance is limited by the availability of biological parameters and validation data. Here, by combining experimental and mathematical models, we have investigated the interactions between cell populations in bone that synchronously orchestrate the bone injury repair program. To do so, we built a mathematical model that captures the dynamics of seven cell populations and the bone mass. Importantly, the system of equations is coupled so that each cell type can regulate the activity of other cell types. This interplay between the different populations poses a challenge regarding the reconciliation between model dynamics and experimental data but gives credence to the novel insights it has allowed us to uncover. These include 1) Anti-inflammatory macrophages drive early osteoclast inhibition and pro-inflammatory phenotype suppression 2) pro-inflammatory macrophages are involved in osteoclast activation (bone resorptive activity), whereas osteoblastic cells promote osteoclast differentiation 3) Pro-inflammatory monocytes/macrophages rise during two expansion waves, which can be explained by the anti-inflammatory macrophages-mediated inhibition phase between the two waves.

Experimentally, as described in our previous study^41^, we observe a rapid expansion of pro-inflammatory monocytes and macrophages in the first 24 hours with anti-inflammatory macrophages emerging shortly thereafter and persisting for up 48 hours. Of the hypotheses tested by the model, *a3-b2-c2* provided the best fit of model simulations to experimental data. Under this set of assumptions, anti-inflammatory macrophages cause retraction of the pro-inflammatory population and facilitates osteoblast expansion and mineralization/stabilization of the injury site. With the natural depletion of the anti-inflammatory population, the remaining injury-associated factors causes a second expansion of pro-inflammatory macrophages and monocytes that in turn enhance osteoclast formation and activity. This increased activity is essential for the resorption of the mineralized callus at the site of injury and the return to bone homeostasis in the given time frame. To our knowledge very few reports have proposed a role and mechanism for this second inflammatory wave during bone healing. A role of MSCs and osteoblastic cells has been proposed for a second increase in inflammatory cytokines like TNF^57^. However, this two waves pattern has been observed in larger spectrum of inflammatory contexts, not only in bone ^52, 108–110^, suggesting that this temporal profile is not bone specific. Importantly, to our knowledge the present study is the first to propose a mathematical framework of bone healing where bone and immune cell populations are fully coupled and to inform such a model with experimental longitudinal data of all these populations.

Our hypothesis combination approach allowed us to explore the polarization properties of monocytes/macrophages that can be difficult to determine *in vivo*. For example, the best fitting ODE model, *a3-b2-c2* allowed us to estimate the rates of pro- and anti-inflammatory macrophage polarization and indicates that pro-inflammatory macrophages do not re-polarize into an anti-inflammatory phenotype given the time frame, which goes against studies suggesting macrophage plasticity and reprogramming, at least in the context of bone injury repair. While not disputing the possibility that macrophages can repolarize, our results suggest that, based on the timing of the acquired experimental timing points, repolarization does not appear to be the main mechanism that recapitulates macrophage polarization dynamics. Additional insights provided by the ODE model include estimations on macrophage lifespan during the healing process and the contributory roles of pro-inflammatory macrophages and monocytes to the process. This information can be critical for therapies that target specific myeloid populations during bone injury repair in a bid to accelerate bone healing.

Another important result of our study is that the hypothesis combination that fit the best, *a3-b2-c2*, implies that osteoblast and pro-inflammatory monocytes-macrophages have distinct roles in osteoclast biology. According to this model, osteoblast drives osteoclast differentiation, whereas pro-inflammatory monocytes-macrophages drive osteoclast resorptive activity. In most studies, this distinction is not made and both cell types are assumed to contribute to both osteoclast differentiation and resorptive activity. The data we present here suggest a distinction in osteoclast supportive functions. This hypothesis combination has been also used to simulate OSM depletion bone injury process. Model simulations were in accordance with experimental data from an independent published study.

An important aspect of mathematical modeling is that it allows us to distill the key cell species and molecules driving the bone dynamics. Of note, our model does not consider the potential roles of other cell types in the bone ecosystem that could contribute, such as T cells. Our results suggest that that integrating myeloid populations into the model provides enough resolution to explain the process of non-critical bone injury repair. Our mathematical framework is flexible enough, however, that the effects of other immune cells such as T cells could be included. Additionally, we are aware the hypotheses we have identified throughout this study, while the most common, do not cover exhaustively all myeloid behaviors described in literature. However, the hypothesis testing pipeline we have devised enables us to efficiently adapt our model to reflect any additional hypotheses. Another important aspect that our model currently neglects is that of bone quality which requires a different set of data acquisition techniques and further refinement of the mathematical approach. Moreover, bone healing is a spatially regulated process and having this aspect included in the model would be an exciting refinement in order to explore further mechanistic aspects of bone structure and regeneration.

Through our hypothesis combination approach, we have integrated established biology into a mathematical framework describing cell population dynamics during non-critical bone injury repair. One potential application of our framework is to investigate how time to healing subsequent to bone injury can be reduced. Existing studies have shown that bone healing times can be impacted in modulating pro- and anti-inflammatory macrophages^13, 111, 112, 119^. While the model is parameterized with mouse data, there is much overlap between mice and humans with respect to the phases of the bone injury repair program. Thus, using our existing workflow, we can conceivably re-parameterize our model with human patient-derived data to further its potential as a relevant prospective tool for the clinic.

In conclusion, we have developed a coupled ordinary differential equation (ODE) system of the bone ecosystem that models the interplay between 8 key cellular populations during bone injury repair. The model yields several novel findings regarding macrophage dynamics and macrophage impact on osteoblasts and osteoclasts dynamics. Further, the model can also provide novel insights into phenomena that are hard to measure *in vivo* such as rate of pro- or anti-inflammatory polarization over time. A better understanding of bone healing will have clinical translatability allowing, for instance, to accelerate the process and improve patient outcomes. The model accounts for coupling between these population and will be useful in developing therapeutic strategies/interventions that shorten healing times. Further, the model has broad applicability and can be used as a platform to examine other bone diseases such as osteoarthritis and skeletal malignancies such as bone-metastatic cancer.

## MATERIALS AND METHODS

### Intratibial Bone Injury Model

Data from in vivo bone injury experiment were derived from preprint manuscript available in the Cold Spring Harbor bioRxiv server, https://www.biorxiv.org/content/10.1101/2020.10.13.338335v1.abstract. In this study, all animal studies were performed in accordance with Guidelines for the Care and Use of Laboratory Animals published by the National Institutes of Health, and approved by the Animal Care and Use Committee at the University of South Florida, under IACUC Protocol R5857 (CCL). Male C57BL/6 mice (5-6 weeks old) were purchased from Jackson Laboratory. Mice (n=30) were subject to tibial bone injury by penetration of a 28-gauge (0.3062mm diameter) syringe through the knee epiphysis to mid-shaft. Mice tibias at baseline and at days 1, 2, 3, 7 and 14 (n=5/time point) were collected for analysis. Temporal population data was used to parameterize subsequent mathematical models.

### Micro-Computed Topography

Bone volume data was derived from formalin-fixed tibias by micro-computed topography (μCT) scanning using Scanco μ35 scanner. Endosteal trabecular bone volume was analyzed 100μm away from the tip of growth plate to clear the dense bone nature of the growth plate. 1000μm along the midshaft of each bone was then scanned and analyzed using built-in functions (n=30 bones; 5/time point).

### TRAcP Staining

After μCT analysis, tibia bones were decalcified using 14% EDTA for 3 weeks for further staining quantitation and analyses. Decalcified bones were sectioned at 4μm thickness. Sections were enzymatically stained for tartrate-resistant acid phosphatase (TRAcP) for osteoclast numbers based on manufacturer’s protocol^122^. Stained slides were imaged using the Evos Auto microscope to capture 20X photos which included injury site and its immediate periphery. All TRAcP positive (red) multinucleated osteoclasts within 5μm radius from injury were counted, and mathematically converted to osteoclasts / bone marrow volume (#OCL/μm^3^) for each bone at each time point.

### Immunofluorescence Staining and Quantitation

FFPE tibia bones were further sequentially sectioned and baked at 56°C in preparation for immunofluorescence staining of osteoblast (RUNX2 at 1:500; Abcam Cat. No. ab81357) and nuclear staining (DAPI). Deparaffined and rehydrated slides were subject to heat-induced antigen retrieval method. Sections were then blocked and incubated in primary antibodies diluted in 10% normal goat serum in TBS overnight at 4°C. Subsequently, slides were stained with secondary Alexa Fluor 568-conjugated antibody at 1:1000 at room temperature for 1 hour under light-proof conditions. Stained slides were stained with DAPI for nuclear contrast and mounted for imaging at 20X using Zeiss upright fluorescent microscope to include the injury site as well as the immediate peripheral tissue. All runx2 positive cells (red staining colocalizing with DAPI) within 5μm radius from injury were counted and mathematically converted to osteoblasts / bone marrow volume (#OBL/μm^3^) for each bone at each time point.

### Flow Cytometry and Analysis

Harvested contralateral injured tibias (n=30; 5/time point) had tips removed and were subjected to centrifugation at 16,000g for 5 seconds for isolation of whole bone marrow for flow cytometry staining and analysis. Red blood cells were lysed using RBC Lysis Buffer from Sigma Aldrich (Cat. No. R7757-100ML) as per manufacturer’s guidelines. Live bone marrow cells were subject to FcR-receptor blocking (1:3; BioLegend; Cat. No. 101319) and viability staining (1:500; BioLegend; Cat. No. 423105). Samples were then stained by cell-surface conjugated antibodies from BioLegend diluted in autoMACS buffer (Miltenyi; Cat. No. 130-091-221) for phenotyping myeloid cells: CD11b-BV786 (1:200; Cat. No. 101243), LY-6C-Alexa Fluor 488 (1:500; Cat. No.128021) and LY-6G-Alexa Fluor 700 (1:200; Cat. No. 561236). Cells were then fixed with 2% paraformaldehyde in dark prior to intracellular staining. Fixed cells were permeabilized using intracellular conjugated antibodies to assess polarization status: NOS2-APC (1:100; eBioscience; Cat. No. 17-5920-80) and ARG1-PE (1:100; R&D; Cat. No. IC5868P). Appropriate compensation and fluorescence-minus-one (FMO) controls were generated in parallel either with aliquots of bone marrow cells or Rainbow Fluorescent Particle beads (BD Biosciences; Cat. No. 556291). All antibody concentrations were titrated prior to injury study using primary bone marrow cells to ensure optimal separation and detection of true negative and positive populations. Stained controls and samples were analyzed using BD Biosciences LSR flow cytometer (Supplemental Figure 2).

## MATHEMATICAL AND COMPUTATIONAL METHODS

### Comprehensive model structure

In order to efficiently describe the process of model building regarding the different hypothesis combinations (Fig. 2), we begin with a generic set of coupled equations, using a formulation that is valid for all hypotheses combinations (Supplemental Fig. 3). For all populations, homeostasis was described by a replenishment term and a clearance term. For polarized monocytes/ macrophages, no replenishment was considered at homeostasis, as the baseline measured by flow cytometry was close to zero. Here is the detailed description, equation by equation:

***Equation 1: Naive Monocytes***

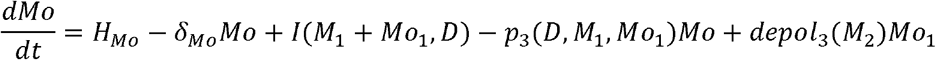

Naive monocytes are assumed to be replenished at a constant rate *H_Mo_*, and to die at a rate *δ_Mo_Mo. δ_Mo_*, the lifespan parameter, was retrieved from literature (Table 3), and *H_Mo_* was estimated so that monocyte level at homeostasis would match experimentally measured monocyte baseline. The term *I*(*M*_1_, *Mo*_1_, *D*) corresponds to the number of monocytes infiltrating the bone marrow per unit of time due to injury factors and pro-inflammatory cells. It is equal to *I*_1_(*M*_1_ + *Mo*_1_) + *I*_2_*D*. As indicated by the mathematical formulation, inflammation-associated monocyte recruitment is driven by injury signals on one hand, and pro-inflammatory monocytes/macrophages on the other hand. This reflects the fact that cellular debris and pro-inflammatory cells produce factors that help recruiting additional monocytes^1^. The term *p*_3_(*D, M*_1_, *Mo*_1_) represents the number of naive monocytes polarizing into pro-inflammatory monocytes per unit of time, as a function of cellular debris and pro-inflammatory cells. The term *depol*_3_(*M*_2_)*Mo*_1_ represents the number pro-inflammatory monocytes reverting back to a naive state per unit of time, as function of anti-inflammatory macrophages.

***Equation 2: Naive Macrophages***

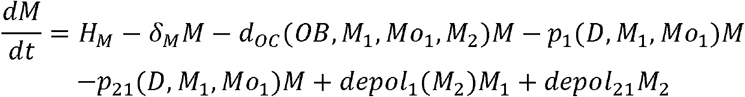

Naive macrophage are assumed to be replenished at a constant rate *H_M_*, and to die at a rate *δ_M_M. δ_M_*, the lifespan parameter, was retrieved from literature (Table 3), and *H_M_* was estimated so that macrophage level at homeostasis would match experimentally measured macrophage baseline. The term *d_oc_*(*OB, M*_1_, *Mo*_1_, *M*_2_)*M* corresponds to the number of macrophages differentiating into osteoclasts per unit of time, as a function of osteoblasts (RANKL, OPG), pro-inflammatory macrophages (IL-1, TNF), pro-inflammatory monocytes (IL-1, TNF) and anti-inflammatory macrophages (IL-10, TGF). The term *p*_1_(*D, M*_1_, *Mo*_1_)*M* represents the number of naive macrophages polarizing into a pro-inflammatory state per unit of time, as a function of injury factors and pro-inflammatory cells. The term *p*_21_(*D, M*_1_, *Mo*_1_)*M* represents the number of naive macrophages polarizing into an anti-inflammatory state per unit of time, as a function of injury factors and pro-inflammatory cells. The term *depol*_1_(*M*_2_)*M*_1_ represents the number of pro-inflammatory macrophages that revert back to a naive state per unit of time, as a function of anti-inflammatory macrophages. The term *depol*_21_*M*_2_ represents the number of anti-inflammatory macrophages that revert back to a naive state per unit of time. Of note, no influx or differentiation from monocytes during inflammation were assumed, as macrophage population does not show evidence of expansion in our obtained biological data.

***Equation 3: Pro-inflammatory Macrophages***

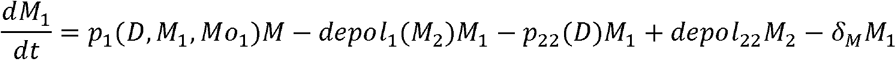

Pro-inflammatory macrophages are assumed to be absent of the bone marrow under homeostatic conditions, as experimental al baseline level did not exceed ?% The term *p*_1_(*D, M*_1_, *Mo*_1_)*M* represents the number of pro-inflammatory macrophages generated (from naive pool) per unit of time, as a function of injury factors and pro-inflammatory cells. The term *depol*_1_(*M*_2_)*M*_1_ represents the number of pro-inflammatory macrophages that revert back to a naive state per unit of time, as a function of anti-inflammatory macrophages. The term *depol*_22_*M*_2_ represents the number of anti-inflammatory macrophages that reprogram into a pro-inflammatory phenotype. The term *p*_22_(*D*)*M*_1_ represents the number of pro-inflammatory macrophages that reprogram into an anti-inflammatory phenotype, as a function of injury factors.

***Equation 4: Anti-inflammatory Macrophages***

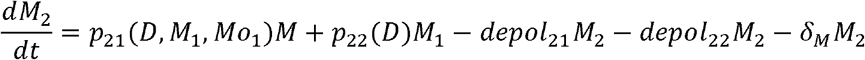

Anti-inflammatory macrophages are assumed to be absent of the bone marrow under homeostatic conditions, as experimental al baseline level did not exceed ?% The term *p*_21_(*D, M*_1_, *Mo*_1_)*M* represents the number of anti-inflammatory macrophages generated (from naive pool) per unit of time, as a function of injury factors and pro-inflammatory cells. The term *p*_22_(*D*)*M*_1_ represents the number of pro-inflammatory macrophages that reprogram into an anti-inflammatory phenotype, as a function of injury factors. The term *depol*_21_*M*_2_ represents the number of pro-inflammatory macrophages that revert back to a naive state per unit of time. The term *depol_22_M_2_* represents the number of anti-inflammatory macrophages that reprogram into a pro-inflammatory phenotype.

***Equation 5: Pro-inflammatory Monocytes***

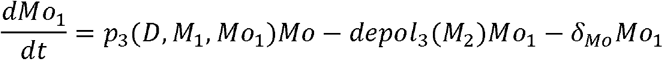

Pro-inflammatory monocytes are assumed to be absent of the bone marrow under homeostatic conditions, as experimental al baseline level did not exceed ?% The term *p*_3_(*D, M*_1_, *Mo*_1_)*Mo* represents the number of pro-inflammatory monocytes generated (from naive pool) per unit of time, as a function of injury factors and pro-inflammatory cells. The term *depol*_3_(*M*_2_)*Mo*_1_ represents the number of pro-inflammatory monocytes that revert back to a naive state per unit of time, as a function of anti-inflammatory macrophages. The term *depol*_22_(*M*_2_) represents the number of anti-inflammatory macrophages that reprogram into a pro-inflammatory phenotype. The term *p*_22_(*D*)*M*_1_ represents the number of pro-inflammatory macrophages that reprogram into an anti-inflammatory phenotype, as a function of injury factors.

### Pro-inflammatory Macrophages/Monocytes and Anti-inflammatory Macrophages: Hypothesis c1

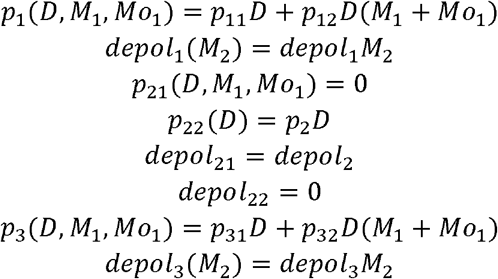

Both CD11b+Ly6C+ monocytes and CD11b+Ly6C-macrophages can polarize into a pro-inflammatory phenotype^1^, in response to injury signals or to factors produced by already present pro-inflammatory cells^1^. In assumptions c1 and c2, monocytes and macrophages polarize into pro-inflammatory monocytes and macrophages respectively through two terms. The first is proportional to the amount of injury signals present and the second is proportional to the amount of pro-inflammatory cells present. In the assumption c3, injury signals polarize local resident macrophages, and those in turn promote polarization of pro-inflammatory monocytes. In c3, pro-inflammatory macrophages repolarize into anti-inflammatory macrophages by the up-take of cellular debris/injury signals^133^. In this scenario, by plasticity, anti-inflammatory macrophages naturally return to a pro-inflammatory phenotype in absence of signals^134^. Polarization into anti-inflammatory macrophages was assumed to be proportional to the amount of injury signals for assumption c1, proportional to the amount of pro-inflammatory cells for assumption c2, and a transition term from pro-inflammatory macrophages for c3.

### Pro-inflammatory Macrophages/Monocytes and Anti-inflammatory Macrophages: Hypothesis c2

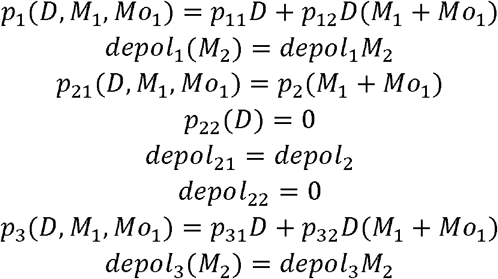

Both CD11b+Ly6C+ monocytes and CD11b+Ly6C-macrophages can polarize into a pro-inflammatory phenotype^1^, in response to injury signals or to factors produced by already present pro-inflammatory cells^1^. In assumptions c1 and c2, monocytes and macrophages polarize into pro-inflammatory monocytes and macrophages respectively through two terms. The first is proportional to the amount of injury signals present and the second is proportional to the amount of pro-inflammatory cells present. In the assumption c3, injury signals polarize local resident macrophages, and those in turn promote polarization of pro-inflammatory monocytes. In c3, pro-inflammatory macrophages repolarize into anti-inflammatory macrophages by the uptake of cellular debris/injury signals^133^. In this scenario, by plasticity, anti-inflammatory macrophages naturally return to a pro-inflammatory phenotype in absence of signals^134^. Polarization into anti-inflammatory macrophages was assumed to be proportional to the amount of injury signals for assumption c1, proportional to the amount of pro-inflammatory cells for assumption c2, and a transition term from pro-inflammatory macrophages for c3.

### Pro-inflammatory Macrophages/Monocytes and Anti-inflammatory Macrophages: Hypothesis c3

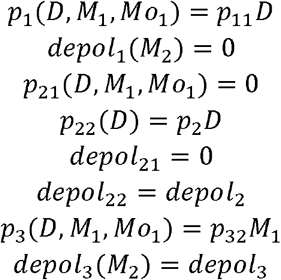

Both CD11b+Ly6C+ monocytes and CD11b+Ly6C-macrophages can polarize into a pro-inflammatory phenotype^1^, in response to injury signals or to factors produced by already present pro-inflammatory cells^1^. In assumptions c1 and c2, monocytes and macrophages polarize into pro-inflammatory monocytes and macrophages respectively through two terms. The first is proportional to the amount of injury signals present and the second is proportional to the amount of pro-inflammatory cells present. In the assumption c3, injury signals polarize local resident macrophages, and those in turn promote polarization of pro-inflammatory monocytes. In c3, pro-inflammatory macrophages repolarize into anti-inflammatory macrophages by the uptake of cellular debris/injury signals^133^. In this scenario, by plasticity, anti-inflammatory macrophages naturally return to a pro-inflammatory phenotype in absence of signals^134^. Polarization into anti-inflammatory macrophages was assumed to be proportional to the amount of injury signals for assumption c1, proportional to the amount of pro-inflammatory cells for assumption c2, and a transition term from pro-inflammatory macrophages for c3.

***Equation 6: Osteoblasts***

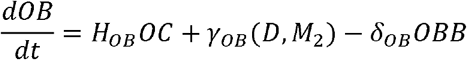

Osteoblast are assumed to be replenished at a rate *H_OB_OC*, proportional to osteoclasts. This reflects the ability of osteoclasts to produce osteogenic signals like transforming growth factor β (TGFβ) and bone morphogenetic proteins (BMPs)^123^. Similar assumptions are considered in published works of homeostatic bone remodeling^33–35^. Osteoblasts are assumed to die at a rate *δ_OB_OBB. δ_OB_*, the lifespan parameter, was retrieved from literature (Table 3), and *H_OB_* was estimated so that osteoblast level at homeostasis would match experimentally measured osteoblast baseline. The term *γ_OB_*(*D, M*_2_) represents the number of osteoblasts generated per unit of time, as a function of injury factors and anti-inflammatory factors.

### Osteoblast dynamics: Hypothesis b1

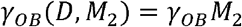

The osteoblast clearance term was assumed to be proportional to the bone volume, in order to account for osteoblast differentiation into osteocytes, when resorbing bone matrix^123^. A similar assumption is made in the model developed by Ryser et al. that describes bone remodeling as a spatial evolutionary game^124^. During injury, an extra term for osteoblast expansion is present, driven by anti-inflammatory macrophages (hypothesis b1) or injury factors (hypothesis b2), both supported by literature^1, 3, 6, 7, 125^.

### Osteoblast dynamics: Hypothesis b2

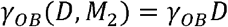

The osteoblast clearance term was assumed to be proportional to the bone volume, in order to account for osteoblast differentiation into osteocytes, when resorbing bone matrix^123^. A similar assumption is made in the model developed by Ryser et al. that describes bone remodeling as a spatial evolutionary game^124^. During injury, an extra term for osteoblast expansion is present, driven by anti-inflammatory macrophages (hypothesis b1) or injury factors (hypothesis b2), both supported by literature^1, 3, 6, 7, 125^.

***Equation 7: Osteoclasts***

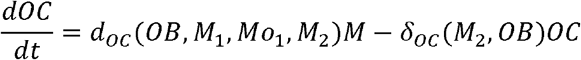

Osteoclasts are assumed to be replenished at a rate *d_OC_*(*OB, M*_1_, *Mo*_1_, *M*_2_)*M*, which reflects differentiation of macrophages into osteoclasts, as a function of osteoblasts, pro-inflammatory macrophages, pro-inflammatory monocytes, anti-inflammatory macrophages^126–128^. This reflects osteoclastic factors produced by osteoblasts (RANKL) and pro-inflammatory monocytes/macrophages (IL-1, TNF), as well as anti-osteoclastic factors produced by osteoblasts (OPG) and anti-inflammatory macrophages (transforming growth factor β (TGFβ), IL-10). The term *δ_OC_*(*M*_2_, *OB*)*OC* represents the number of osteoclasts dying per unit of time, as a function of anti-inflammatory macrophages and osteoblasts. This reflects factors produced by anti-inflammatory macrophages (IL-10, TGFβ) and osteoblasts (OPG) that reduce osteoclast lifespan.

### Osteoclast dynamics: Hypothesis a1

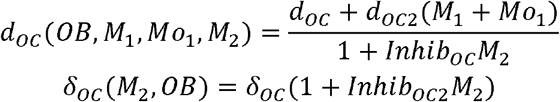

This osteoclast formation term was assumed to be proportional to osteoblasts for assumption a3, reflecting the ability of osteoblastic cells to produce RANKL, which is an essential mediator of osteoclast formation^123^. For the other assumptions, homeostatic osteoclast replenishment was assumed to be constant. This term had an additional contribution from pro-inflammatory monocytes/macrophages for assumptions a1 and a2, representing the ability of pro-inflammatory monocytes/macrophages to produce factors like IL-1 and TNF that favor osteoclast formation^123, 129^. Osteoclast formation was divided by an inhibitory term, a linear function of anti-inflammatory macrophages for assumptions a1 and a3, and a linear function of osteoblasts for assumption a2. The first assumption reflects factors produced by anti-inflammatory macrophages, like IL-10, that disrupt osteoclast formation^88^, whereas the second reflects the ability of osteoblasts to produce osteoprotegerin (OPG), a RANKL decoy receptor^123^. Moreover, this inhibition affects not only the ability of monocytes-macrophages to fuse and form osteoclasts, but also their life span. Indeed RANKL is necessary for osteoclast survival since OPG produced by osteoblasts reduces their life span^130^. Similarly, anti-inflammatory macrophages produce TGFβ, which is known to drive osteoclast apoptosis^131^.

### Osteoclast dynamics: Hypothesis a2

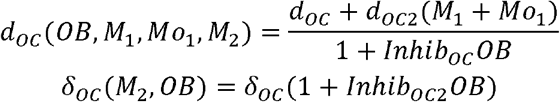

This osteoclast formation term was assumed to be proportional to osteoblasts for assumption a3, reflecting the ability of osteoblastic cells to produce RANKL, which is an essential mediator of osteoclast formation^123^. For the other assumptions, homeostatic osteoclast replenishment was assumed to be constant. This term had an additional contribution from pro-inflammatory monocytes/macrophages for assumptions a1 and a2, representing the ability of pro-inflammatory monocytes/macrophages to produce factors like IL-1 and TNF that favor osteoclast formation^123, 129^. Osteoclast formation was divided by an inhibitory term, a linear function of anti-inflammatory macrophages for assumptions a1 and a3, and a linear function of osteoblasts for assumption a2. The first assumption reflects factors produced by anti-inflammatory macrophages, like IL-10, that disrupt osteoclast formation^88^, whereas the second reflects the ability of osteoblasts to produce osteoprotegerin (OPG), a RANKL decoy receptor^123^. Moreover, this inhibition affects not only the ability of monocytes-macrophages to fuse and form osteoclasts, but also their life span. Indeed RANKL is necessary for osteoclast survival since OPG produced by osteoblasts reduces their life span^130^. Similarly, anti-inflammatory macrophages produce TGFβ, which is known to drive osteoclast apoptosis^131^.

### Osteoclast dynamics: Hypothesis a3

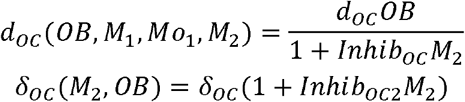

This osteoclast formation term was assumed to be proportional to osteoblasts for assumption a3, reflecting the ability of osteoblastic cells to produce RANKL, which is an essential mediator of osteoclast formation^123^. For the other assumptions, homeostatic osteoclast replenishment was assumed to be constant. This term had an additional contribution from pro-inflammatory monocytes/macrophages for assumptions a1 and a2, representing the ability of pro-inflammatory monocytes/macrophages to produce factors like IL-1 and TNF that favor osteoclast formation^123, 129^. Osteoclast formation was divided by an inhibitory term, a linear function of anti-inflammatory macrophages for assumptions a1 and a3, and a linear function of osteoblasts for assumption a2. The first assumption reflects factors produced by anti-inflammatory macrophages, like IL-10, that disrupt osteoclast formation^88^, whereas the second reflects the ability of osteoblasts to produce osteoprotegerin (OPG), a RANKL decoy receptor^123^. Moreover, this inhibition affects not only the ability of monocytes-macrophages to fuse and form osteoclasts, but also their life span. Indeed RANKL is necessary for osteoclast survival since OPG produced by osteoblasts reduces their life span^130^. Similarly, anti-inflammatory macrophages produce TGFβ, which is known to drive osteoclast apoptosis^131^.

***Equation 8: Bone Volume***

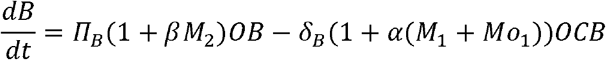

Bone dynamics is described by two terms: a bone resorption term, which is the volume of bone resorbed per unit of time and is assumed to be proportional to the number of osteoclasts, and a bone formation term, which is the volume of bone formed per unit of time and is assumed to be proportional to the number of osteoblasts. Such assumptions have been broadly used across a large variety of modeling studies^33–35^. *δ_B_*(1 + *α*)*OCB* is the bone resorption term and is the sum of the two terms *δ_B_OCB* (homeostatic resorption) and *α*(*M*_1_ + *Mo*_1_)*δ_B_OCB* (pro-inflammatory monocyte/macrophage mediated resorption). As indicated by this mathematical formulation, bone resorption was assumed to also be proportional to the bone mass. This reflects the fact that more bone volume increases the likelihood for bone resorption. Furthermore, this formulation ensures bone mass stays strictly positive in the model. *π_B_*(1 + *β*)*OB* is the bone formation term and is the sum of the two terms *π_B_OB* (homeostatic bone formation) and *βM*_2_*π_B_OB* (anti-inflammatory macrophage mediated bone formation). Under homeostasis, bone is formed at rate *π_B_OB* and resorbed at rate *δ_B_OCB*. Resorption rate *δ_B_*, as well as as parameters *α*. and *β*, were calibrated on bone volume dynamics, and bone formation rate *π_B_* was then imposed by the relation *π_B_OB*_0_ = *δ_B_OC*_0_*B*_0_, which ensures that bone volume remains at homeostasis when osteoblasts and osteoclasts are at homeostatic levels. As indicated by mathematical formulations, bone formation and resorption terms were assumed to linearly increase with respect to anti-inflammatory and pro-inflammatory monocytes/ macrophages, respectively. This accounts for the fact that anti-inflammatory macrophages typically produce osteogenic factors like TGFβ or OSM, that are known to promote osteoblast expansion and bone mineralization^3^, and for the fact that pro-inflammatory monocytes/macrophages typically produce osteolytic factors like TNF and IL-1, that are known to promote osteoclast resorptive activity^132^.

***Equation 9: Injury Factors***

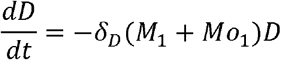

Injury factors dynamics consists in an exponential type of decay, with a decay rate *δ*(*M*_1_ + *Mo*_1_) proportional to pro-inflammatory monocytes/macrophages number, which represents how pro-inflammatory monocytes/macrophages uptake cellular debris, which in return reduces pro-inflammatory signals.

### Population homeostasis

In order to estimate the homeostatic cell replenishment parameters, we set them equal to the clearance term (lifespan) which was either based on literature values or calibrated directly from experimental data (Supplemental Fig. 1, Supplemental Tables 1 and 4).

### ODE solver

The ODE45 function of Matlab was used to solve the differential equation system. The experimental baseline values (time 0) were used as initial conditions.

### Parameter estimation method

To estimate parameters for goodness of fit, we defined the following objective function:

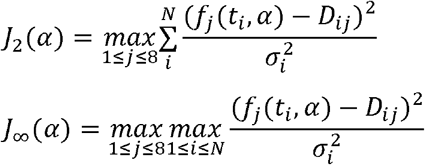

Where i represents the time point index and j the variable index, □ represents the parameter set used to evaluate the model function f, *D_ij_* represents the experimental data of variable j at time point i, *σ_i_* represents the experimental standard deviation (over all the animals of a given time point), and N represents the number of time points. The functional *J*_2_ corresponds to the weighted least squares criterion. The functional *J*_∞_ is the Tchebychev approximation, wich considers the maximal residual instead of the sum of the residuals. In both cases, the choice of the max over the observed variables of the sum of the squares of the residuals was motivated to make sure that every variable was fitted with equal relative importance. Indeed, in the case of the minimization of the sum of the squares over all the variables, it is sometimes possible to find an optimum in minimizing certain variables at the detriment of others. This way, we ensure that all variables are equally well fitted.

The reason for considering the objective function *J*_∞_ in addition to the classical criterion *J*_2_ is to avoid neglecting any time point in the fit. The least squares functional allows sometimes to find an optimum optimizing certain time points at the detriment of others. In this current study, this is a potentially big issue, as the time sampling is not homogenous across the time points, meaning that biological dynamics of importance might be ignored, while still producing a good fit under the least squares metric.

In order to minimize this function representing the error estimate between data and model, we used the Matlab function fminsearch with a penalization term to stay in a parameter range set with reasonable boundaries.

In order to rank the models in term of goodness of fit, we used AIC, that is defined as follows:

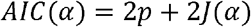

where p is the number of parameters, and the functional J is either *J*_2_ or *J_∞_*.

### OSM knockout data

In order to retrieve data from the OSM knockout independent dataset, we used webplotdigitizer to collect datapoint from the plot presented in Guihard et al^50^.

## Authors’ Contributions

C. H. Lo, E. Baratchart C. C. Lynch and D. Basanta were responsible for study conception and design. C. H. Lo, E. Baratchart C. C. Lynch and D. Basanta developed the methodology. C. H. Lo and E. Baratchart acquired the data while C. H. Lo, E. Baratchart C. C. Lynch and D. Basanta were responsible for analysis and interpretation of data. C. H. Lo, E. Baratchart C. C. Lynch and D. Basanta were responsible for manuscript writing and editing.

---

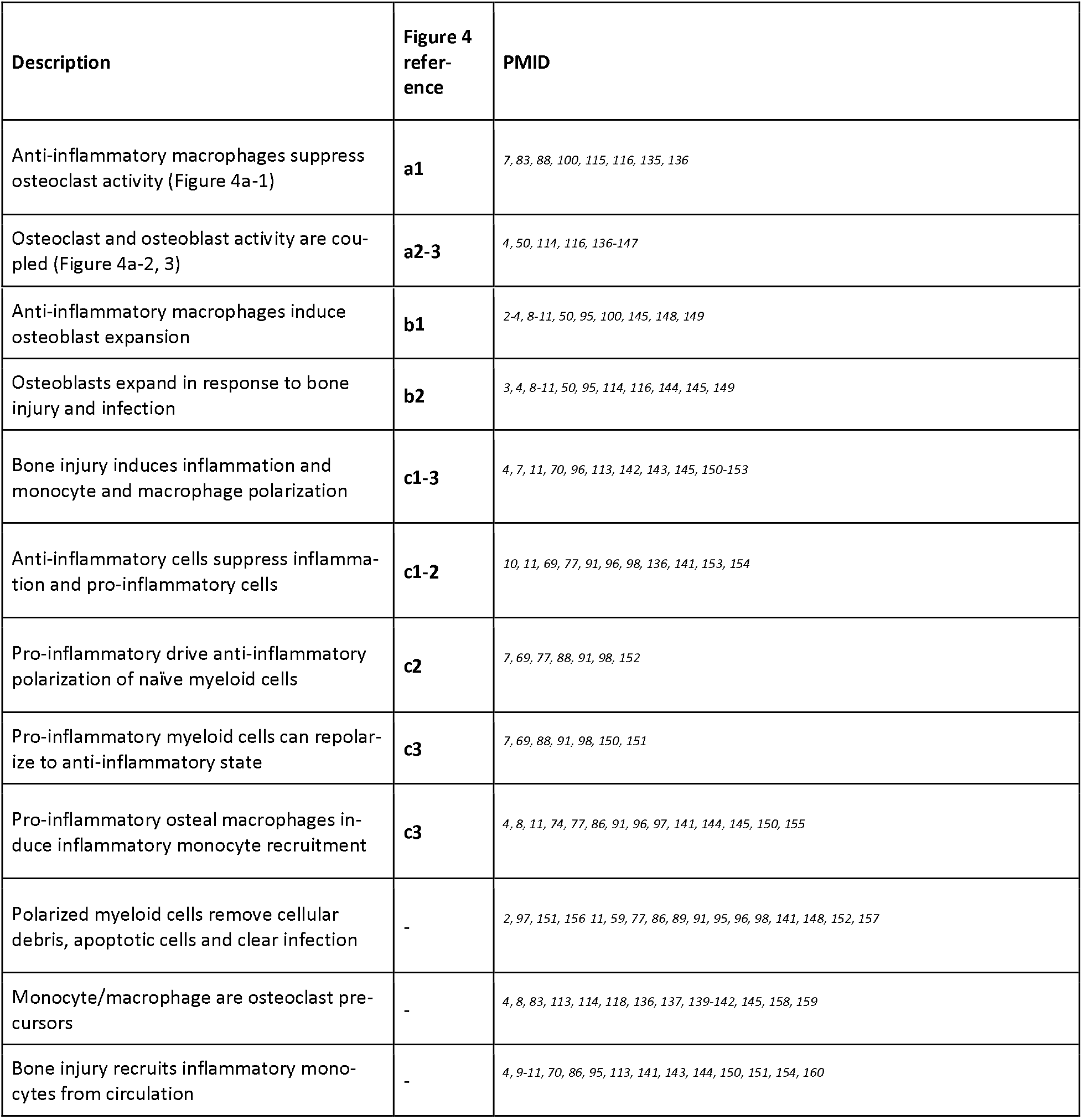

## SUPPLEMENTARY FIGURE LEGENDS

**Supplemental Fig. 1**. ODE equations for the best fitting model: Hypothesis combination a3b2c2 The ODE systems govern behaviors of each population and are parameterized by published values when available, such as the natural lifespan of monocytes (*δ_Mo_* in *dMo/dt*). Parameters with no reference publication were estimated to obtain best possible fits to temporal dynamics data (parameters in red) and are listed in Table 3. In all equations, black terms correspond to homeostatic dynamics, whereas red terms correspond to injury dynamics.

**Supplemental Fig. 2**. Alternative hypotheses combination a3 b1 c2 (green boxes in **a-c**) computational results. **a** Mechanism a3 assumes that osteoblasts and anti-inflammatory macrophages promote and inhibit osteoclast formation, respectively. **b** Mechanism b1 assumes that anti-inflammatory macrophages promote osteoblast expansion. **c** Mechanism c2 assumes that injury factors promote pro-inflammatory monocytes/macrophages polarization. Pro-inflammatory monocytes/macrophages promote anti-inflammatory macrophages polarization, which in return drive depolarization of pro-inflammatory monocytes/macrophages back to the naive state. **d** Schematic representation of the model using a3b1c2 hypothesis combination. Arrows represent positive (green) or negative (red) types of cellular interactions. **e** Temporal plots and corresponding goodness of fit metrics (AIC and R2s) across all populations, obtained through *J*_∞_ minimization.

**Supplemental Fig. 3**. Generic formulation of coupled ODE models, valid for all hypotheses combinations. Each term (e.g formation rate, clearance, transition,…) is a functional form of other variables reflecting cellular interactions described in Figure 2. Black terms correspond to homeostasis, red terms correspond to injury dynamics and are described in details in Mathematical methods.

**Supplemental Fig. 4**. Alternative hypotheses combination a3 b2 c1 (green boxes in **a-c**) computational results. **a** Mechanism a3 assumes that osteoblasts and anti-inflammatory macrophages promote and inhibit osteoclast formation, respectively. **b** Mechanism b2 assumes that injury factors promote osteoblast expansion. **c** Mechanism c1 assumes that injury factors promote pro-inflammatory monocytes/macrophages and anti-inflammatory macrophages polarization. Anti-inflammatory macrophages drive depolarization of pro-inflammatory monocytes/macrophages back to the naive state. **d** Schematic representation of the model using a3b2c1 hypothesis combination. Arrows represent positive (green) or negative (red) types of cellular interactions. **e** Temporal plots and corresponding goodness of fit metrics (AIC and R2s) across all populations, obtained through *J*_∞_ minimization.

**Supplemental Fig. 5**. Alternative hypotheses combination a3 b1 c1 (green boxes in **a-c**) computational results. **a** Mechanism a3 assumes that osteoblasts and anti-inflammatory macrophages promote and inhibit osteoclast formation, respectively. **b** Mechanism b21assumes that injury factors promote osteoblast expansion. **c** Mechanism c1 assumes that anti-inflammatory macrophages promote pro-inflammatory monocytes/macrophages and anti-inflammatory macrophages polarization. Anti-inflammatory macrophages drive depolarization of pro-inflammatory monocytes/macrophages back to the naive state. **d** Schematic representation of the model using a3b2c1 hypothesis combination. Arrows represent positive (green) or negative (red) types of cellular interactions. **e** Temporal plots and corresponding goodness of fit metrics (AIC and R2s) across all populations, obtained through *J*_∞_ minimization.

**Supplemental Fig. 6**. Alternative hypotheses combination a2 b1 c1 (green boxes in **a-c**) produces the second worst fit of all hypotheses combinations. **a** Mechanism a2 assumes that pro-inflammatory and macrophages and osteoblasts promote and inhibit osteoclast formation, respectively. **b** Mechanism b1 assumes that anti-inflammatory macrophages promote osteoblast expansion. **c** Mechanism c1 assumes that injury factors promote pro-inflammatory monocytes/macrophages polarization and anti-inflammatory macrophages. The latter drive depolarization of pro-inflammatory monocytes/macrophages back to the naive state. **d** Schematic representation of the model using a2b1c1 hypothesis combination. Arrows represent positive (green) or negative (red) types of cellular interactions. **e** Temporal plots and corresponding goodness of fit metrics (AIC and R2s) across all populations, obtained through *J*_∞_ minimization.

**Table 1**. Established biological behaviors and functions of bone cell populations. Framework for a comprehensive and coupled 9-population ODE model is constructed based off of summarizing known published interactions between each population. Inclusion of select hypotheses for each ambiguous aspect of myeloid biology is based on the prevalence of their corresponding publications (at least seven supporting references for each).

**Table 2**. Akaike information criterion (AIC) for comprehensive ODE of all 18 combinations of hypotheses. Left columns denote the hypotheses from each of three mechanisms tested. The AIC scores resulting from *J*_2_ and *J*_∞_ minimization for each model are shown on the right and vary dramatically across models. Comparing AICs reveal one best combination (boxed in red) and the worst fitting model is highlighted in blue lines.

**Table 3**. Estimated parameters of the two best-fitting models. First column is the parameter notation used in the equations. Second column is the biological meaning of the parameter. Third and fourth columns are the parameter values for both models. Fifth column is the parameter unit. Sixth column is the reference used for retrieving parameter value, when it was possible/available. Parameters for which no reported estimation could be found were estimated by fitting on experimental data.

**Table 4**. Table showing biological description of each mathematical variable with data-derived initial conditions and units. First column is the variable notation used in the equations for each parameter. Second column is the biological meaning of the variable. Third column is the initial condition for each variable, typically an initial cell population level. Fourth column is the variable unit.

**Table 5**. Residuals lower than one for Mathematical model of all 18 combinations of hypotheses, resulting from *J*_2_ and *J*_∞_ minimization. For each hypothesis combination, the table shows how many residuals are lesser than 1 over all 40 residuals, which equates how many times the model lies within the experimental error bar.

