## Supplementary material for "Integrated computational and *in vivo* models reveal Key Insights into Macrophage Behavior during bone healing": Tables

**Table 1.** Reference sources of predominant mechanisms

| Description | Figure 4 reference | Reference |
|---|---|---|
| Anti-inflammatory macrophages suppress osteoclast activity | **a1** | *7, 83, 88, 100, 115, 116, 135, 136* |
| Osteoclast and osteoblast activity are coupled | **a2-3** | *4, 50, 114, 116, 136-147* |
| Anti-inflammatory macrophages induce osteoblast expansion | **b1** | *2-4, 8-11, 50, 95, 100, 145, 148, 149* |
| Osteoblasts expand in response to bone injury and infection | **b2** | *3, 4, 8-11, 50, 95, 114, 116, 144, 145, 149* |
| Bone injury induces inflammation and monocyte and macrophage polarization | **c1-3** | *4, 7, 11, 70, 96, 113, 142, 143, 145, 150-153* |
| Anti-inflammatory cells suppress inflammation and pro-inflammatory cells | **c1-2** | *10, 11, 69, 77, 91, 96, 98, 136, 141, 153, 154* |
| Pro-inflammatory drive anti-inflammatory polarization of naïve myeloid cells | **c2** | *7, 69, 77, 88, 91, 98, 152* |
| Pro-inflammatory myeloid cells can repolarize to anti-inflammatory state | **c3** | *7, 69, 88, 91, 98, 150, 151* |
| Pro-inflammatory osteal macrophages induce inflammatory monocyte recruitment | **c3** | *4, 8, 11, 74, 77, 86, 91, 96, 97, 141, 144, 145, 150, 155* |
| Polarized myeloid cells remove cellular debris, apoptotic cells and clear infection | - | *2, 97, 151, 156, 11, 59, 77, 86, 89, 91, 95, 96, 98, 141, 148, 152, 157* |
| Monocyte/macrophage are osteoclast precursors | - | *4, 8, 83, 113, 114, 118, 136, 137, 139-142, 145, 158, 159* |
| Bone injury recruits inflammatory monocytes from circulation | - | *4, 9-11, 70, 86, 95, 113, 141, 143, 144, 150, 151, 154, 160* |

**Table 2.** Akaike information criterion (AIC) for comprehensive ODE of all 18 combinations of hypotheses

| Mechanism Hypothesis | | | $J_2$ | $J_\infty$ |
|---|---|---|---|---|
| **a** | **b** | **c** | **AIC Score** | **AIC Score** |
| *1* | *1* | *1* | 79 | 73 |
| *1* | *2* | *3* | 77 | 68 |
| *1* | *1* | *3* | 78 | 67 |
| *1* | *2* | *1* | 79 | 73 |
| *1* | *1* | *2* | 89 | 70 |
| *1* | *2* | *2* | 115 | 65 |
| *2* | *2* | *1* | 81 | 76 |
| *2* | *1* | *1* | 103 | 79 |
| *2* | *1* | *2* | 78 | 69 |
| *2* | *1* | *3* | 78 | 66 |
| *2* | *2* | *2* | 88 | 78 |
| *2* | *2* | *3* | 128 | 113 |
| *3* | *1* | *1* | 58 | 45 |
| *3* | *1* | *3* | 76 | 56 |
| *3* | *2* | *1* | 57 | 46 |
| *3* | *2* | *3* | 86 | 59 |
| *3* | *1* | *2* | 51 | 42 |
| ***3*** | ***2*** | ***2*** | **44** | **39** |

**Table 3.** Parameter values of the winning models

| Parameter | Description | Value<br>*a3 b2 c2* | Unit | Reference |
|---|---|---|---|---|
| $\delta_{Mo}$ | Monocyte Lifespan | 0.45 | $Day^{-1}$ | *179* |
| $\delta_{M}$ | Macrophage Lifespan | 0.1 | $Day^{-1}$ | *108* |
| $\delta_{OB}$ | Bone-mediated Osteoblast lifespan | 0.32 | $Day^{-1}$ | Estimated |
| $\delta_{OC}$ | Osteoclast lifespan | 0.53 | $Day^{-1}$ | *176* |
| $\gamma_{OB}$ | Macrophage-mediated Osteoblast Formation Rate | $4.33 \times 10^{4}$ | Cell $mm^{-3}$ $Day^{-1}$ | Estimated |
| $\delta_{B}$ | Per Bone Volume Unit Homeostatic Resorption Rate | $5.99 \times 10^{-7}$ | $Cell^{-1}$ $Day^{-1}$ | Estimated |
| $\Pi_{B}$ | Homeostatic Bone Apposition Rate | $6.018 \times 10^{-7}$ | $mm^{3}$ $Cell^{-1}$ $Day^{-1}$ | Determined from $\delta_{B}$ |
| $\alpha$ | Modulation of Resorption Rate by Pro-inflammatory Cell | 0.0022 | $Cell^{-1}$ | Estimated |
| $\beta$ | Modulation of Apposition Rate by Anti-inflammatory Cell | 0.012 | $Cell^{-1}$ | Estimated |
| $H_{Mo}$ | Homeostatic Monocyte Formation Rate | $1.5 \times 10^{4}$ | Cell $Day^{-1}$ | Estimated |
| $H_{M}$ | Homeostatic Macrophage Formation Rate | $1.8 \times 10^{4}$ | Cell $Day^{-1}$ | Estimated |
| $H_{OB}$ | Homeostatic Osteoblast Formation Rate | 0.014 | Cell $Cell^{-1}$ $Day^{-1}$ | Determined from $\delta_{OB}$ |
| $d_{OC}$ | Osteoblast-mediated Osteoclast Formation Rate | $5.35 \times 10^{-5}$ | $Cell^{-1}$ $Day^{-1}$ | Determined from $\delta_{OC}$ |
| $Inhib_{OC}$ | Macrophage-mediated Osteoclast inhibition | 0.016 | $Cell^{-1}$ | Estimated |
| $Inhib_{OC2}$ | Macrophage-mediated Osteoclast inhibition | 0.052 | $Cell^{-1}$ | Estimated |
| $\delta_{D}$ | Macrophage/Monocyte-mediated debris clearance | $1.71 \times 10^{-5}$ | $Cell^{-1}$ $Day^{-1}$ | Estimated |
| $I_{1}$ | Pro-Inflammatory cells-mediated monocyte recruitment | $1.21 \times 10^{-22}$ | Cell $Cell^{-1}$ $Day^{-1}$ | Estimated |
| $I_{2}$ | Injury signals-mediated monocyte recruitment | $6.15 \times 10^{3}$ | Cell $mm^{-3}$ $Day^{-1}$ | Estimated |
| $p_{31}$ | Injury Factors-mediated Pro-inflammatory Monocytes Polarization Rate | $6.094 \times 10^{-73}$ | $mm^{-3}$ $Day^{-1}$ | Estimated |
| $p_{32}$ | Pro-inflammatory Cells-mediated Pro-inflammatory Monocytes Polarization Rate | $3.42 \times 10^{-5}$ | $mm^{-3}$ $Cell^{-1}$ $Day^{-1}$ | Estimated |
| $depol_{3}$ | Pro-inflammatory Monocytes depolarization Rate | 0.029 | $Cell^{-1}$ $Day^{-1}$ | Estimated |
| $p_{11}$ | Injury Factors-mediated Pro-inflammatory M Polarization Rate | $5.86 \times 10^{-9}$ | $mm^{-3}$ $Day^{-1}$ | Estimated |
| $p_{12}$ | Pro-inflammatory Cells-mediated Pro-inflammatory Macrophages Polarization Rate | $4.69 \times 10^{-4}$ | $mm^{-3}$ $Cell^{-1}$ $Day^{-1}$ | Estimated |
| $p_{2}$ | Anti-Inflammatory Macrophages Polarization Rate | $2.34 \times 10^{-6}$ | $Cell^{-1}$ $Day^{-1}$ | Estimated |
| $depol_{1}$ | Pro-inflammatory Macrophages depolarization Rate | 0.37 | $Cell^{-1}$ $Day^{-1}$ | Estimated |
| $depol_{2}$ | Anti-inflammatory Macrophages depolarization Rate | $1.47 \times 10^{-39}$ | $Day^{-1}$ | Estimated |

**Table 4.** Model variables description

| Mathematical variable | Biological variable | Initial conditions | Units |
|---|---|---|---|
| OB | Osteoblasts | 3.1E+04 | Cell number |
| OC | Osteoclasts | 8.8E+04 | Cell number |
| B | Bone | 0.3530 | $mm^3$ |
| D | Cellular debris/Injury factors | 2.8 | $mm^3$ |
| Mo | Naive monocytes | 1.4E+04 | Cell number |
| M | Naive macrophages | 2.8E+04 | Cell number |
| M1 | Pro-inflammatory macrophages | 0 | Cell number |
| M2 | Anti-inflammatory macrophages | 0 | Cell number |
| Mo1 | Pro-inflammatory monocytes | 0 | Cell number |

**Table 5.** Residuals lower than one for comprehensive ODE of all 18 combinations of hypotheses

| Mechanism Hypothesis | | | $J_2$ | $J_\infty$ |
|---|---|---|---|---|
| **a** | **b** | **c** | **Residulas<1** | **Residulas<1** |
| *1* | *1* | *1* | 18/40 | 14/40 |
| *1* | *2* | *3* | 16/40 | 12/40 |
| *1* | *1* | *3* | 20/40 | 15/40 |
| *1* | *2* | *1* | 15/40 | 14/40 |
| *1* | *1* | *2* | 12/40 | 14/40 |
| *1* | *2* | *2* | 9/40 | 13/40 |
| *2* | *2* | *1* | 18/40 | 11/40 |
| *2* | *1* | *1* | 15/40 | 12/40 |
| *2* | *1* | *2* | 18/40 | 17/40 |
| *2* | *1* | *3* | 20/40 | 15/40 |
| *2* | *2* | *2* | 16/40 | 15/40 |
| *2* | *2* | *3* | 8/40 | 8/40 |
| *3* | *1* | *1* | 16/40 | 13/40 |
| *3* | *1* | *3* | 13/40 | 14/40 |
| *3* | *2* | *1* | 17/40 | 14/40 |
| *3* | *2* | *3* | 16/40 | 19/40 |
| *3* | *1* | *2* | 21/40 | 15/40 |
| ***3*** | ***2*** | ***2*** | **27/40** | **25/40** |
